# Profound Treg perturbations correlate with COVID-19 severity

**DOI:** 10.1101/2020.12.11.416180

**Authors:** Silvia Galván-Peña, Juliette Leon, Kaitavjeet Chowdhary, Daniel A. Michelson, Brinda Vijaykumar, Liang Yang, Angela Magnuson, Zachary Manickas-Hill, Alicja Piechocka-Trocha, Daniel P. Worrall, Kathryn E. Hall, Musie Ghebremichael, Bruce D. Walker, Jonathan Z. Li, Xu G. Yu, MGH COVID-19 Collection & Processing Team, Diane Mathis, Christophe Benoist

**Author notes:** Address correspondence to: Christophe Benoist, Department of Immunology, Harvard Medical School, 77 Avenue Louis Pasteur, Boston, MA 02115. Equal contribution (Silvia Galván-Peña, Juliette Leon).

## Abstract

The hallmark of severe COVID-19 disease has been an uncontrolled inflammatory response, resulting from poorly understood immunological dysfunction. We explored the hypothesis that perturbations in FoxP3+ T regulatory cells (Treg), key enforcers of immune homeostasis, contribute to COVID-19 pathology. Cytometric and transcriptomic profiling revealed a distinct Treg phenotype in severe COVID-19 patients, with an increase in both Treg proportions and intracellular levels of the lineage-defining transcription factor FoxP3, which correlated with poor outcomes. Accordingly, these Tregs over-expressed a range of suppressive effectors, but also pro-inflammatory molecules like IL32. Most strikingly, they acquired similarity to tumor-infiltrating Tregs, known to suppress local anti-tumor responses. These traits were most marked in acute patients with severe disease, but persisted somewhat in convalescent patients. These results suggest that Tregs may play nefarious roles in COVID-19, via suppressing anti-viral T cell responses during the severe phase of the disease, and/or via a direct pro-inflammatory role.

## INTRODUCTION

COVID-19 resulting from SARS-CoV2 infection is a major global health challenge. Many infected individuals remain asymptomatic or present with only mild flu-like symptomatology, appearing 5-10 days after exposure, and clearing over 1-2 weeks. But this is followed, in patients with more severe disease, by immunopathology and immune dysregulation of poorly understood origin, a major root of the fatality rate of 1-12 % in different locales. Broad immune-profiling studies (Mathew et al., 2020; Laing et al., 2020; Su et al., 2020) have documented several immune phenomena that track with disease severity: lymphopenia (Huang et al., 2020), myeloid cell abnormalities (Schulte-Schrepping et al., 2020), impaired response to interferon (Hadjadj et al., 2020), and high levels of inflammatory cytokines (“cytokine storm”) (Del Valle et al., 2020). Multiomic studies have shown that these manifestations are embedded within multi-trait immunotypes (Mathew et al., 2020; Laing et al., 2020; Su et al., 2020), complicating mechanistic inference and the definition of potential therapies.

Regulatory T cells (Tregs) expressing the transcription factor FoxP3 are essential to maintain immunologic homeostasis, self-tolerance, and to prevent runaway immune responses (Josefowicz et al., 2012). Tregs regulate the activation of several lineages of the innate and adaptive immune systems, through several effector mechanisms (Vignali et al., 2008). Furthermore, particular populations of “tissue Tregs” provide homeostatic regulation in several non-immunological tissues, controlling inflammation and promoting harmonious tissue repair (Panduro et al., 2016). On the other hand, Tregs can also prove noxious, as evidenced most clearly by their suppression of effective cytotoxic responses in tumors, situations in which they adopt a distinctive phenotype (De Simone et al., 2016; Plitas et al., 2016; Magnuson et al., 2018). They can also have paradoxical effects on antiviral responses (Lund et al., 2008; Almanan et al., 2017)

In light of these contrasting influences, we hypothesized that Tregs might contribute to the balance of disease manifestations that distinguish mild from severe outcomes after SARS-CoV2 infection: for instance by insufficiently curtailing the inflammatory component, by over-curtailing the anti-viral response, or by phenotypic destabilization. We thus performed a deep immunologic and transcriptional analysis of circulating blood Tregs across a cohort of confirmed COVID-19 patients.

## RESULTS

### More Tregs, more FoxP3 in severe COVID-19 patients

We thus hypothesized that Tregs might contribute to COVID-19 pathology, and performed a deep immunologic and transcriptional analysis of circulating blood Tregs across a cohort of confirmed COVID-19 patients (n=57, Fig. 1A, Table S1; mild: outpatients; severe: hospitalized, 65% of which in intensive care (ICU), mostly sampled during the cytokine storm period; recovered: virus-negative convalescents). Flow cytometry, with a multiparameter panel that parsed CD25^+^ FoxP3^+^ Tregs and their different phenotypes (gating strategies in Fig. S1A), revealed several perturbations (representative plots in Fig. 1B). First, several severe patients showed increased Treg proportions among CD4+ T cells; for some as an increased proportion only (likely resulting from preferential resistance among CD4+ T cells during lymphopenia); for others, from true numerical increase relative to healthy donors (HD) (Fig. 1C; this conclusion differs from a recent analysis of CD4+ T cells that up-regulated CD69/4.1BB after 24hrs’ culture with a multi-peptide cocktail, hence very different from the present *ex vivo* study and possibly complicated by selection in culture (Meckiff et al., 2020)). Recovered patients largely reverted to baseline. Second, expression of both FoxP3 and CD25 varied in severe patients, bidirectionally for CD25 (Fig. S1B), but as a reproducible increase of FoxP3 mean fluorescence intensity (MFI; Fig. 1D). No hint of FoxP3 induction was observed in conventional CD4^+^ T cells (Tconv). Increased FoxP3 expression coincided with the increase in Treg percentage in most but not all patients (Fig. S1C), and was observed within a broad window of 30-50 days after symptom onset, including some recently recovered patients (Fig. 1E). It was not related to body/mass index (BMI), a risk factor for COVID-19 (Fig. S1D), but coincided with disease severity, particularly marked in ICU-admitted patients (Fig. S1E), and in lymphopenic cases (Fig. S1F). No dependence of these phenotypes was found on any particular treatment, in particular glucocorticoids (Table S1). FoxP3 expression was not directly correlated with blood levels of the inflammatory marker C-reactive protein (CRP), but essentially all FoxP3^hi^ patients had elevated CRP (Fig. S1G). Therefore, severe COVID-19 entails a striking induction of FoxP3 expression in Tregs, in a manner not been previously observed in any autoimmune or infectious context.

**Fig 1.**
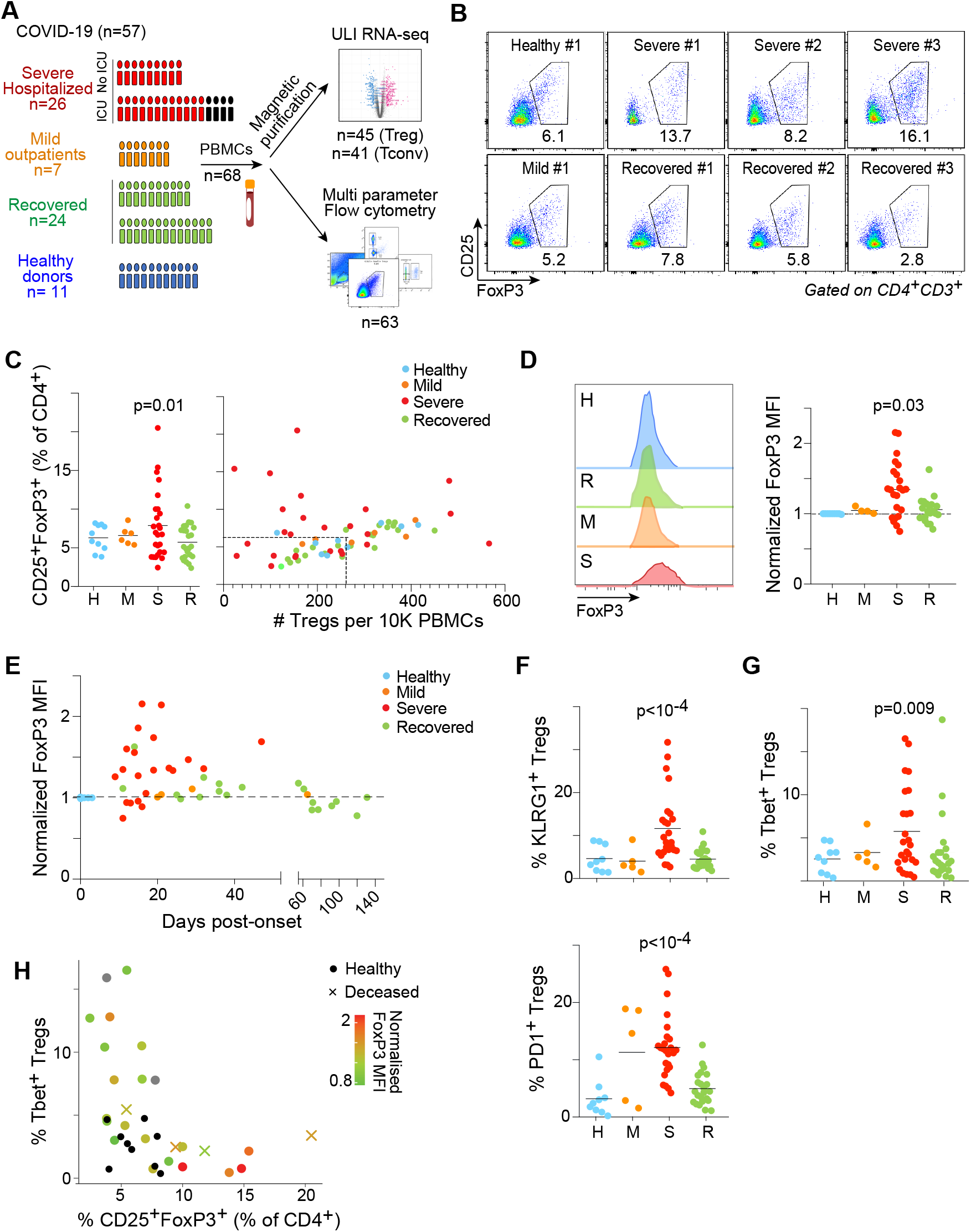
Treg over-representation and FoxP3 induction in COVID-19 patients. (**A**)Experimental approach. Tregs from PBMCs across mild, severe and recovered COVID-19 patients, compared to healthy donors, were assessed by flow cytometry, as well as by RNA-seq. (**B**) Representative flow cytometry plots of CD25^+^ FoxP3^+^ Tregs from COVID-19 patients’ PBMCs. (**C**) Proportions (left) and proportions vs absolute numbers (right) of Tregs as measured by flow cytometry across donors; H: HD; M: Mild; S: Severe; R: Recovered. p.values from random permutation test quantitating number of outlier values relative to distribution in HDs. (**D**) FoxP3 expression, measured as MFI in CD127^lo^ CD25^+^ Tregs. Representative flow cytometry profiles (left) and quantification (right); Mann–Whitney p.values (**E**) Correlation between FoxP3 expression and days post disease symptoms onset across COVID-19 patients (**F**) Proportion of KLRG1+ (left) and PD1+ (right) Tregs as determined by flow cytometry across COVID-19 patients; significance computed as for C. (**G**) Proportion of Tbet+ Tregs as determined by flow cytometry across COVID-19 patients; significance computed as for C. (**H**) Correlation between percentage of CD25^+^ FoxP3^+^ Tregs (x-axis), percentage of Tbet+ Tregs (y-axis) and FoxP3 expression as MFI (color gradient) within severe COVID-19 patients. Healthy controls depicted in black dots and patients with fatal outcome by a cross.

We examined the expression of several markers and transcription factors (TF) to further assess Treg evolution during COVID-19. CD45RA, which marks naïve Tregs, was reduced in patients, especially in those with heightened Tregs and FoxP3 (Fig. S1H,I). Expression of the activation markers KLRG1 and PD1 was increased (Fig. 1F), although not necessarily coordinately (Fig. S1J). Suppression of specific T cell effector functions is associated with the expression in Tregs of the same key TFs that drive those Tconv functions (Josefowicz et al., 2012). No patient sample showed significant expression of Bcl6, which marks T follicular regulators (Linterman et al., 2011; Chung et al., 2011). But Tbet, expressed in Tregs that preferentially control Th1 responses, was over-represented in severe patient Tregs (Fig. 1G). Interestingly, severe patients with frequent Tbet+ Tregs were distinguished from those with highest Treg expansion and FoxP3 over-expression (Fig. 1H). Ultimately deceased patients were mostly found in the latter group. Thus, severe COVID-19 seems to elicit divergent deviations among Treg cells. We surmise that the presence of Tbet^+^ Tregs is related to the control of Th1 antiviral response among effector cells.

### Accentuated Treg transcriptome in severe COVID-19 patients

To decipher the functional consequences of FoxP3 overexpression in Tregs from severe COVID-19 patients, we generated 86 RNAseq transcriptome profiles, passing quality thresholds, of blood Treg (CD4^+^ CD25^hi^ CD127^lo^) or Tconv (CD4^+^ CD25^-^). Donors largely coincided with those analyzed above, and we opted to profile purified populations rather than single-cell (higher throughput, lower processing/computational costs) after magnetic purification (Fig. S2A). The datasets matched profiles from flow-purified Tregs, with the usual differential expression of “Treg signature” genes (Fig. S2B). In line with the cytometry, Tregs from severe COVID-19 patients showed higher *FOXP3* expression (Fig. S2C). This over-expression had a consequence: Tregs from severe COVID-19 patients displayed a heightened expression of Treg signature transcripts (Fig. 2A), reflected by a high “TregSignature Index”, most markedly biased in the severe group, but also in mild and recovered patients (Fig. 2B), and correlating with *FOXP3* expression (Fig. S2C). This bias was not homogeneous across all Treg-up signature genes, a ranked plot of differential expression in each donor revealing a small subset of TregUp signature genes actually downregulated in severe COVID-19 patients (including *TLR5, ID3, FCRL1*; Fig. 2C). At the other end of the spectrum, several Treg effector or activation transcripts were up-regulated in severe patients (*ENTPD1, HPGD, IL12RB2*). Most transcripts encoding known Treg effector molecules showed were up-regulated, albeit to various degrees (Fig. 2D), with the marked exception of *AREG,* a dominant player in Treg promotion of tissue repair (Burzyn et al., 2013; Arpaia et al., 2015). We also examined transcripts associated with effectiveness at suppressing different Tconv functions (Josefowicz et al., 2012). Transcripts associated with T follicular regulators were largely unaffected in Tregs from severe patients (Fig. 2E), with the exception of *PDCD1* (encodes PD1), consistent with cytometry results. In contrast, they generally up-regulated markers related to Th1 suppression (*CXCR3, GZMK, IL12RB1* or *TBX21* (encodes Tbet)), especially marked in patients with low percentage of Tregs (Fig. 2E), in line with the over-representation of Tbet+ Tregs noted above. Profiles from Tconv cells did not denote a particular bias towards any Th phenotype (Fig. S2D,E). Thus, Treg traits observed in the flow cytometry data were confirmed by the transcriptomic signature of these Tregs, which tends towards a super-suppressive phenotype in severe COVID-19 patients.

**Fig 2.**
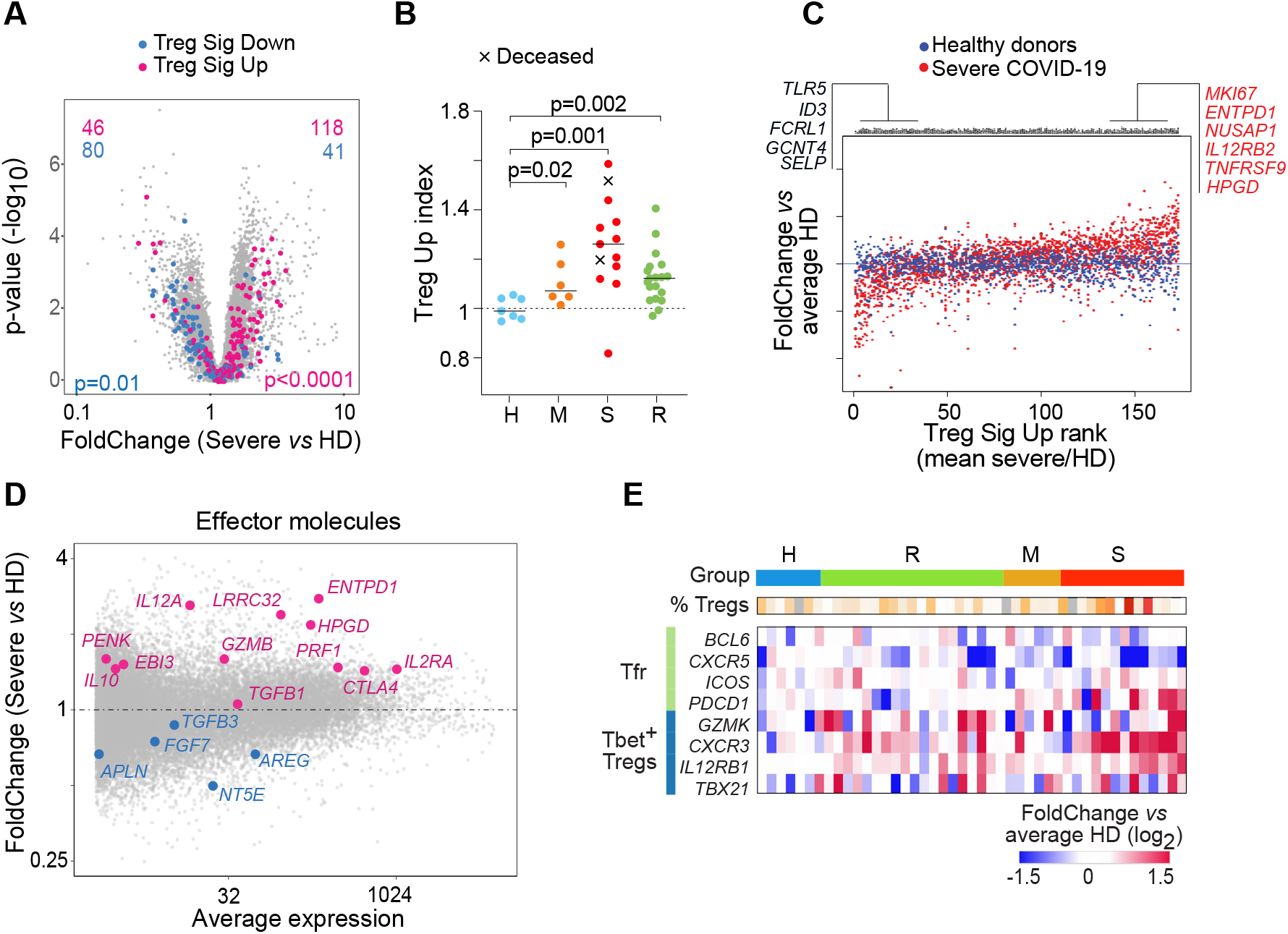
A “Super-Treg” identity in Tregs from severe COVID-19 patients. Tregs from COVID-19 patients and healthy donors (HD) were magnetically purified for population RNA-Seq. (**A**) FoldChange vs p value (volcano) plot of gene expression in Tregs from severe COVID-19 patients compared to HD. Genes from TregUp signature (red) and TregDown signature (blue) are highlighted (Ferraro et al., 2014); p-values from Fisher’s exact. (**B**) Treg-Up index was computed by averaged normalized expression of all signature genes in Tregs from COVID-19 patients and HD. Patients with fatal outcomes depicted as a cross; p-values from Mann–Whitney test (**C**) Ranked expression of Treg-Up signature genes in Tregs from severe COVID-19 patients (red) and HD. The y-axis corresponds to the expression in each donor Treg relative to the mean of HD Tregs. Genes are ranked by their average ratio in severe COVID-19 patients. Each dot is one gene in one donor Treg sample. (**D**) FoldChange vs average expression plot from severe COVID-19 patient Tregs compared to HD. Upregulated (red) and downregulated (blue) Treg effector molecule transcripts highlighted. (**E**) Heatmap of the expression of transcripts typical of T follicular regulators (Tfr) and Tbet+ Treg in Tregs from each donor (as FoldChange vs mean expression in HD). One column per donor, with severity groups color-coded and percentage of FoxP3^+^ CD25^+^ Tregs from the flow cytometry indicated.

### A distinct COVID-19 disease signature Tregs correlates with severity

We then asked more generally what changes, beyond Treg signature and effector transcripts, characterized Tregs in severe COVID-19 patients, relative to HD (Fig. 3A, Table S2; hereafter ‘Severe COVID19 Treg Signature’, SCTS); only a minority of these transcripts belong to the classic Treg signature analyzed above. A SCTS Index computed from this geneset was high in all severe patients, but also persisted in muted fashion in many recovered patients (Fig. 3B), correlating with *FOXP3* expression (Fig. S3A). The index was not directly related to patient age or BMI (Fig. S3B). Grouping SCTS transcripts into a biclustered heatmap (Fig. 3C) revealed several interesting features. Most donors clustered according to severity group, in relation to disease duration (more marked deviation at shorter times), but not to donor age (except inasmuch as severe patients were generally older). Transcripts could be parsed into 8 different modules of distinct composition. Among these, module M4 was almost exclusively composed of cell cycle-related transcripts, and strongly correlated with FoxP3 MFI, indicating that Tregs in COVID-19 patients are highly proliferative, plausibly a compensation for the lymphopenia in many severe patients. M2 mostly included Interferon-stimulated genes (ISGs). The other co-regulated modules included transcripts related to immune cell crosstalk (Fig. 3C, right). Expression of these modules evolved differently during the course of the disease (Fig. 3D). The ISGs in M2 were mostly over-represented at early times, consistent with sharp induction at the early phase of anti-viral responses. Changes in other modules, especially M3 and M6, were shared across all disease stages, indicating a broader and longer-lasting perturbation of the Treg pool, likely caused by secondary consequences of the disease rather than by the virus itself. Focusing on cytokines produced by Tregs, *IL10* was only modestly induced, but *CD70* (CD27L) and *IL32* dominated (Fig. 3E). The latter was intriguing in this context of the cytokine storm that occurs in these severe patients, since it is mainly a pro-inflammatory mediator, in positive feedback with IL6 and IL1β (Zhou and Zhu, 2015). To answer the common question (whether transcriptional changes reflect a shift in subset balance or all shared by all Tregs), we re-analyzed a single-cell RNAseq PBMC dataset from COVID-19 patients (Wilk et al., 2020), drilling down on identifiable Treg cells (Fig. S3C). Tregs from severe patients were generally shifted in the UMAP projection relative to HD (Fig. 3F), and displayed an up-regulation of the SCTS (Fig. 3G, Fig. S3D). *IL32* was again one of the dominant up-regulated transcripts (Fig. S3E).

**Figure 3.**
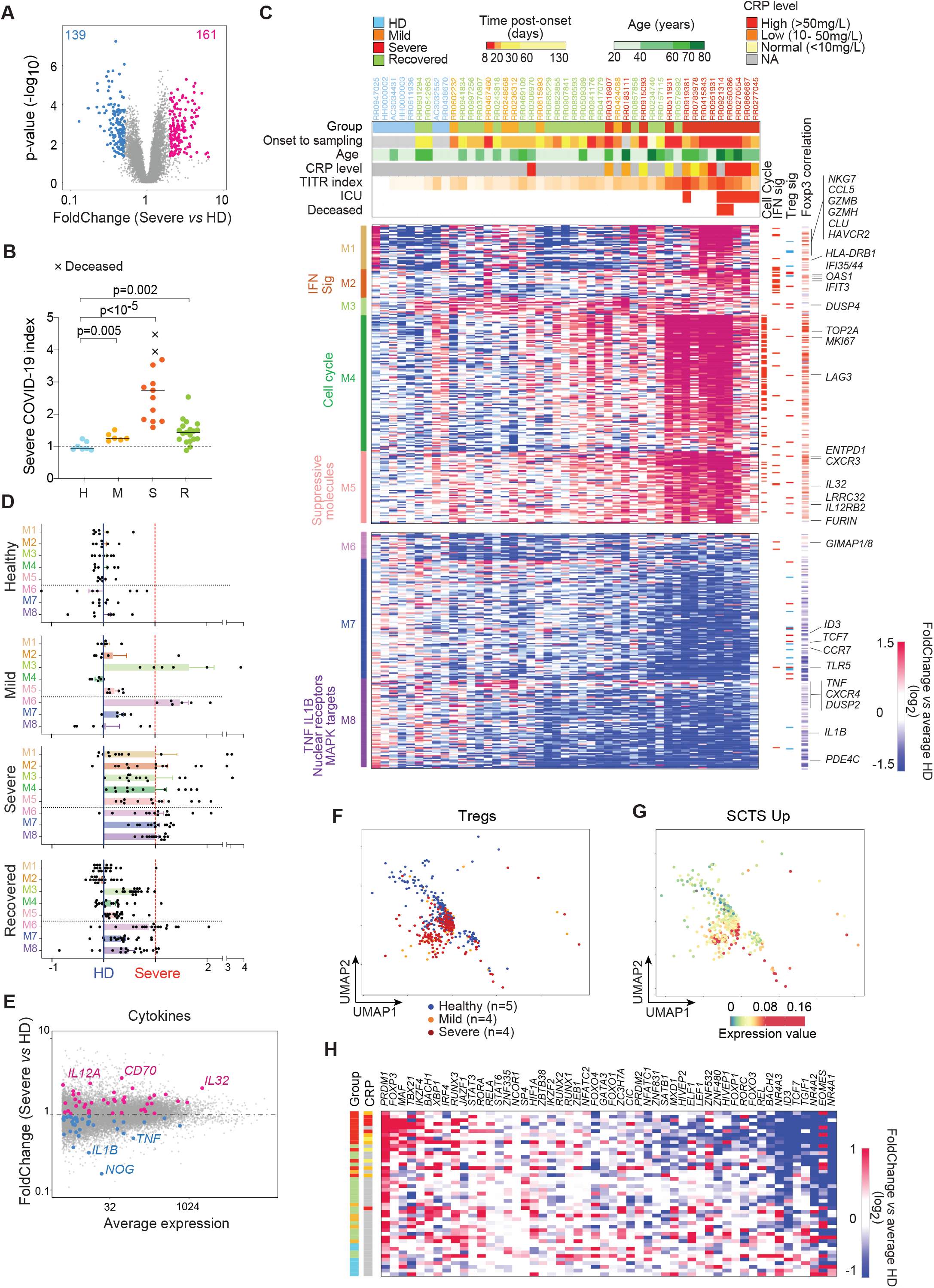
Broad perturbations of Treg transcriptomes in severe COVID-19. (**A**) FoldChange vs p value plot of gene expression in Tregs from severe COVID-19 patients compared to HD. Differentially expressed genes are highlighted (at an arbitrary threshold of p<0.05, FC >2 or <0.5). (**B**) Severe COVID-19 index (computed from relative expression of selected genes from A) in Tregs from COVID-19 patients and HD; p-values from Mann–Whitney test. (**C**) Clustered heatmap of differentially expressed genes (selected as p<0.05, FC >2 or <0.5) across all donors (as ratio to mean of HD values). Each column represents one donor. Top ribbons indicate for each individual: severity group, days from symptom onset to sample collection, age, CRP level at sampling, tumor infiltrating Treg index, ICU admission and final outcome (deceased in red). Left ribbon indicates the coregulated modules and their dominant composition; right ribbons denote transcripts related to cell cycle, Interferon responsive genes, Treg signature genes, and Pearson correlation of each gene’s expression to FOXP3 MFI across all samples. (**D**) Representation of the average expression of each module across each group (from C, mean and SEM). Score computed independently for each module, where 0 corresponds to the average expression of the module in HD Tregs and 1 the average expression in severe COVID-19 Tregs (red line). (**E**) Changes in expression of cytokine-encoding transcripts (on a FoldChange vs average expression plot, severe COVID-19 versus HD Tregs). (**F**) Treg cells extracted from scRNAseq dataset (GSE150728) displayed as a 2D UMAP. The samples are color-coded by group. (**G**) Same plot as F, where each cell is color-coded according to expression of the SCTS-Up signature genes. (**H**) Expression of selected transcription factors in Tregs from each donor (as ratio to mean of HD values). Each column corresponds to one individual, with severity group color-coded and a ribbon indicating CRP levels.

### COVID-19 Tregs as tumor Tregs?

We then attempted to better understand the origin of the SCTS. Comparing changes in Treg and Tconv cells showed some induction in the latter (particularly for the inflammation-related components of M1), but overall much less than in Tregs (Fig. S3F), indicating that the SCTS is largely Treg-specific. With regard to the cytokines, the upregulation of *IL10* and *IL32* were specific to Tregs, as was the down regulation of inflammatory cytokines (*TNF, IL1B*) (Fig S2E). Interestingly, the SCTS was associated with changes in several TFs previously associated with differential gene expression in activated Tregs (Fig. 3H). *PRDM1* (aka BLIMP1) and *MAF* were upregulated, while *ID3*, *TCF7* and *BACH2* were repressed, consistent with previous reports (Cretney et al., 2011; Xu et al., 2018; Maruyama et al., 2011; van Loosdregt et al., 2013; Grant et al., 2020). More unexpected was the strong downregulation of *NR4A1*, normally an indicator of TCR signaling, which may indicate a decoupling of TCR-delivered signals.

Gene enrichment analyses using a curated database of transcriptome variation in CD4+ T cells brought forth 229 datasets with significant overlap to the SCTS (Fig. S4A). Besides the expected interferon-related genesets related to M2, many were related to Tconv and Treg activation, mostly overlapping with M4 and M5. Most intriguing were overlaps with datasets from tumor-infiltrating Tregs. Indeed, the expression of a signature that distinguishes colorectal tumor-infiltrating Tregs (TITR) from normal colon tissue (Magnuson et al., 2018) was strikingly biased in Tregs from severe COVID-19 patients (Fig. 4A). The same bias was found with genesets that distinguish breast and lung TITRs from blood Tregs (De Simone et al., 2016; Plitas et al., 2016) (Fig. 4B-C). Computing a “TITR index” from the colorectal tumor geneset showed that biased expression of tumor Treg transcripts was present in all severe patients, particularly in those eventually deceased (Fig. 4D). This index remained slightly perturbed after recovery, and was highly correlated with the SCTS (Fig. S4B). Conversely, with the exception of M1 and M6, all SCTS modules showed biased expression in tumor Tregs relative to normal tissue (Fig. S4C), indicating widespread sharing not solely limited to activation-related transcripts.

**Figure 4.**
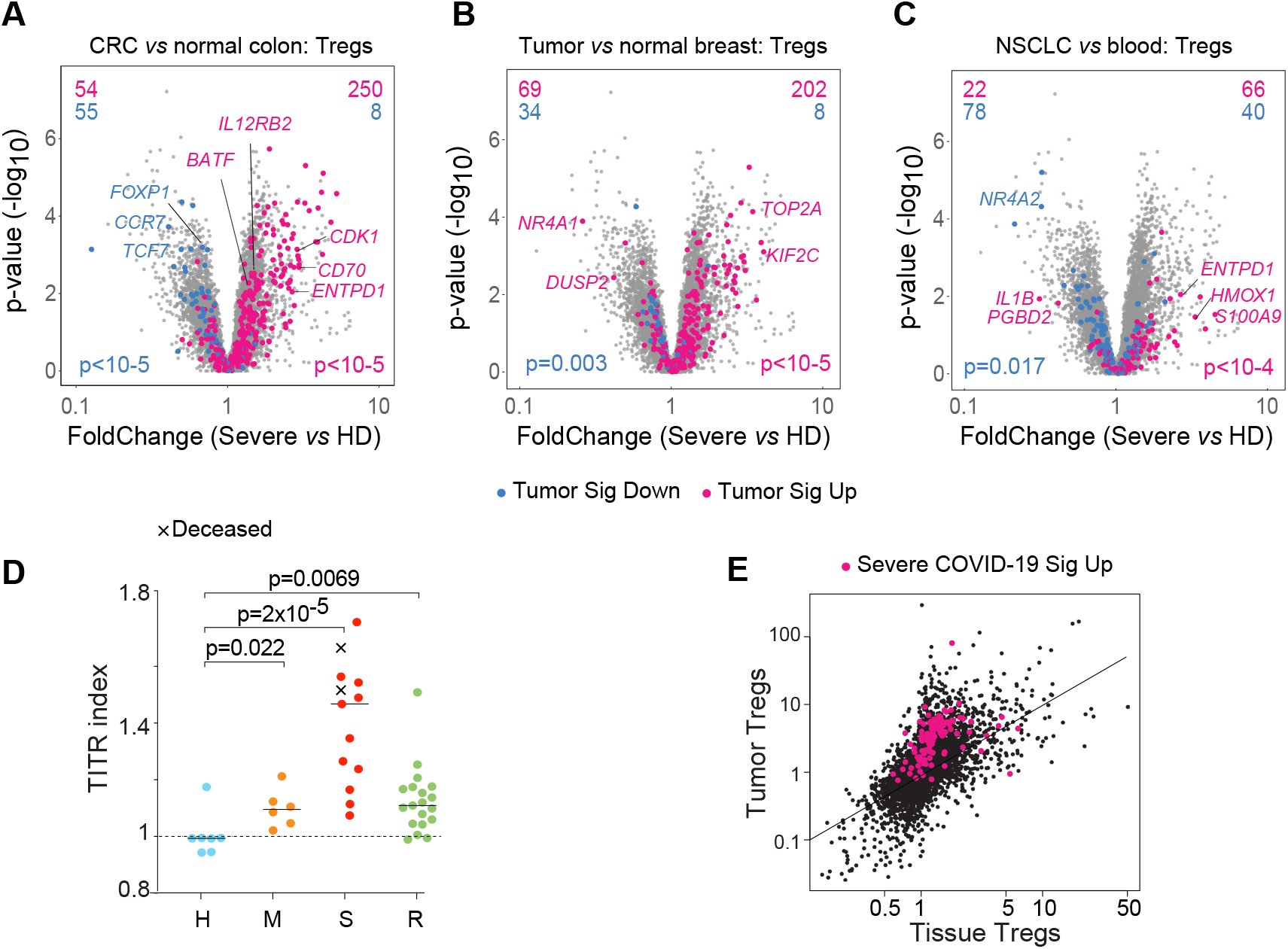
The severe COVID-19 Treg transcriptome overlaps with that of tumor-infiltrating Tregs. (**A – C**) Volcano plots comparing Tregs from severe COVID-19 patients relative to healthy donors (HD), highlighted with signature genes from **A:**colorectal cancer (CRC) vs colon Tregs (GSE116347); **B**: breast tumors vs normal breast Tregs (GSE89225); **C**: non-small-cell lung cancer (NSLC) vs blood Tregs (PRJEB11844); p-values from Fisher’s exact. (**D**) Tumor infiltrating Treg index in Tregs from COVID-19 patients and HD; p-values from Mann Whitney test. (**E**) Highlight of the severe SCTS Up signature (red) on a plot comparing transcriptome shifts in TITRs vs tissue-resident Tregs (see (Magnuson et al., 2018)).

TITRs and tissue Tregs are transcriptionally similar, suggesting the possibility that the SCTS simply denoted Tregs circulating en masse from the lung tissue. This was not the case, however: when highlighted in a direct comparison of tissue and tumor Tregs (from (Magnuson et al., 2018)), the SCTC was clearly biased towards the tumor angle (Fig. 4E). Further, a set of transcripts shared by tissue Tregs showed little enrichment in Tregs from severe COVID-19 patients (Fig. S4D). This bidirectional matching shows that COVID-19 disease appears to be turning blood Tregs into some equivalent of tumor Tregs.

## DISCUSSION

The discovery of this striking Treg phenotype in severe COVD-19 patients, with a unique up-regulation of FoxP3 expression and high transcriptional similarity to tumor Tregs, raises two key questions. First, how do they arise? They are not virally infected (no viral RNA reads to speak of in these cells), and none of the therapies administered to these patients correlates with the Treg traits. Rather, the phenotype must be induced by the immunologic milieu in these patients, and this uniquely in Tregs since Tconvs are far less branded. T cell receptor-mediated stimulation seems unlikely, given the widespread effect on Tregs in the single-cell data which likely transcends clonotypic specificity, and the strong loss of Nur77 (*NR4A1*). A non-specific trigger seems more likely, such as cytokines, interferon, DAMPs released by dying cells, or even more speculatively signals from non-structural viral proteins. Interferon might be involved, and ISGs are part of the signature. We note the strong representation of TNF Receptor family members among the induced SCTS transcripts. Also consistently upregulated is *CXCR3*, the receptor for CXCL10, one of the soluble mediators most induced in severe COVID-19 blood (Filbin et al., 2020). Finally, one factor is common to the tumor microenvironment and to severe COVID-19, hypoxia, which is known to promote Treg suppressive function (Facciabene et al., 2011). Thus, a combination of factors may conspire to achieve the Treg phenotype.

Second, do these aberrant Tregs contribute to COVID-19 physiopathology? Patients with fewer Tregs, lower FoxP3 and less intense SCTS do better, raising the usual issue of inferring causality from correlation. On one hand, these Tregs might be beneficial, controlling a cytokine storm that would have been worse without their unusual contribution. On the other hand, their over-expression of FoxP3, of Treg effector molecules, and their similarity with dominantly suppressive tumor Tregs suggest that COVID-19 Tregs may overly dampen the anti-viral response during the cytokine storm phase (all severe patients profiled were from that period), contributing to the secondary re-expansion of disease. In addition, the intriguing up-regulation of the pro-inflammatory cytokine IL32 (Zhou and Zhu, 2015) might mark these Tregs as amplifying the cytokine storm instead of controlling it. There are precedents for rogue Tregs which acquire pro-inflammatory characteristics (Overacre-Delgoffe et al., 2017) – although admittedly in opposite conditions of FoxP3 attrition.

In summary, adding a key element to the multifaceted COVID-19 immune response, we identify a unique Treg deviation in COVID-19 patients, which might impact on COVID-19 pathogeny as it does in tumors.

## ACKNOWLEGMENTS

We thank Dr C. Blish discussion and access to unpublished data; F. Chen for help with sample processing, and the Broad Genetic Platform for profiling in difficult times. This work was funded by grants from the Massachusetts Consortium for Pathogen Readiness (MassCPR), RO1 AI150686 to CB, and R24 AI072073 to the ImmGen consortium. The MGH/MassCPR COVID biorepository was supported by a gift from Ms. Enid Schwartz, by the Mark and Lisa Schwartz Foundation, the MassCPR and the Ragon Institute of MGH, MIT and Harvard. SG-P was supported by a fellowship from the European Molecular Biology Organisation (ALTF 547-2019), JL by an INSERM Poste d’Accueil and an Arthur Sachs scholarship, KC and DAM by NIGMS-T32GM007753 and a Harvard Stem Cell Institute MD/PhD Training Fellowship, MG by NIH/NIAID P30-AI 060354 to HU-CFAR.

## COLLABORATORS

### MGH COVID-19 Collection & Processing Team participants

#### Collection Team

Kendall Lavin-Parsons, Blair Parry, Brendan Lilley, Carl Lodenstein, Brenna McKaig, Nicole Charland, Hargun Khanna, Justin Margolin

***Department of Emergency Medicine, Massachusetts General Hospital, Boston, MA 02115, USA***.

#### Processing Team

Anna Gonye, Irena Gushterova, Tom Lasalle, Nihaarika Sharma

***Massachusetts General Hospital Cancer Center, Boston, MA 02115, USA***.

Brian C. Russo, Maricarmen Rojas-Lopez

***Division of Infectious Diseases, Department of Medicine, Massachusetts General Hospital, Boston, MA 02115, USA***.

Moshe Sade-Feldman, Kasidet Manakongtreecheep, Jessica Tantivit, Molly Fisher Thomas

***Massachusetts General Hospital Center for Immunology and Inflammatory Diseases, Boston, MA 02115, USA***.

#### Massachusetts Consortium on Pathogen Readiness

Betelihem A. Abayneh, Patrick Allen, Diane Antille, Katrina Armstrong, Siobhan Boyce, Joan Braley, Karen Branch, Katherine Broderick, Julia Carney, Andrew Chan, Susan Davidson, Michael Dougan, David Drew, Ashley Elliman, Keith Flaherty, Jeanne Flannery, Pamela Forde, Elise Gettings, Amanda Griffin, Sheila Grimmel, Kathleen Grinke, Kathryn Hall, Meg Healy, Deborah Henault, Grace Holland, Chantal Kayitesi, Vlasta LaValle, Yuting Lu, Sarah Luthern, Jordan Marchewka (Schneider), Brittani Martino, Roseann McNamara, Christian Nambu, Susan Nelson, Marjorie Noone, Christine Ommerborn, Lois Chris Pacheco, Nicole Phan, Falisha A. Porto, Edward Ryan, Kathleen Selleck, Sue Slaughenhaupt, Kimberly Smith Sheppard, Elizabeth Suschana, Vivine Wilson

***Massachusetts General Hospital, Boston, MA, USA***.

Galit Alter, Alejandro Balazs, Julia Bals, Max Barbash, Yannic Bartsch, Julie Boucau, Josh Chevalier, Fatema Chowdhury, Kevin Einkauf, Jon Fallon, Liz Fedirko, Kelsey Finn, Pilar Garcia-Broncano, Ciputra Hartana, Chenyang Jiang, Paulina Kaplonek, Marshall Karpell, Evan C. Lam, Kristina Lefteri, Xiaodong Lian, Mathias Lichterfeld, Daniel Lingwood, Hang Liu, Jinqing Liu, Natasha Ly, Ashlin Michell, Ilan Millstrom, Noah Miranda, Claire O’Callaghan, Matthew Osborn, Shiv Pillai, Yelizaveta Rassadkina, Alexandra Reissis, Francis Ruzicka, Kyra Seiger, Libera Sessa, Christianne Sharr, Sally Shin, Nishant Singh, Weiwei Sun, Xiaoming Sun, Hannah Ticheli, Alex Zhu

***Ragon Institute, MGH, MIT and Harvard, Cambridge, MA, USA***.

George Daley, David Golan, Howard Heller, David Knipe, Arlene Sharpe

***Harvard Medical School, Boston, MA, USA***.

Nikolaus Jilg, Alex Rosenthal, Colline Wong

***Brigham and Women’s Hospital, Boston, MA, USA***

## MATERIALS AND METHODS

### Patients samples and clinical data collection

Peripheral blood samples from 57 adults infected with SARS-CoV-2 (defined by the concomitant presence of a concordant symptomatology and a positive SARS-CoV-2 real-time reverse-transcriptase–polymerase-chain-reaction (RT-PCR) using nasopharyngeal swabs) prospectively collected in Massachusetts General Hospital (MGH, Boston, USA), were included in this study (Table S1A and S1B). Samples from 11 adult healthy donors (HD) were also collected, dated before December 2019 or defined as no prior diagnosis of or recent symptoms consistent with COVID-19 and a negative PCR test within 3 days pre-collection. COVID-19 patients were split into three different groups depending on the disease severity or the time-course post infection. Patients were categorized as “severe” if admitted to the hospital for moderate to severe COVID-19 symptoms, either in floor or in intensive care unit, as “mild” group if they were outpatients, as “recovered” if they presented a prior positive COVID-19 PCR test, but the most current PCR test was negative. For inpatients, clinical data, including oxygenation status, medications and the outcome of the hospitalization (discharge or death), were recorded from the electronic medical record. Laboratory results were abstracted from the collection date, or if unavailable the closest to that of research blood collection, mostly within 48h. Clinical and demographic information including age, BMI and whether patients were on immunosuppressant or anti-inflammatory drugs are summarized across the groups and displayed individually, respectively in Table S1A and B. All participants provided written informed consent in accordance with protocols approved by the Partners Institutional Review Board and the MGH Human Subjects Institutional Review Board. De-identified Treg analysis was performed per approved HMS-IRB protocol 15-0504-04.

#### Peripheral blood mononuclear cells (PBMC) isolation

5 to 10 mL of whole blood was collected in K2 EDTA tubes and processed within a few hours. An equal volume of buffer (2 mM EDTA in PBS) and blood was mixed and carefully layered over 5ml Ficoll Hypaque solution (GE Healthcare). After centrifugation for 20 min at 900 g (with no break) at 25°C, the mononuclear cell layer was washed twice with excess buffer (three times the volume of the mononuclear cells layer), and centrifuged for 5 min at 400 g. To remove platelets, the cell suspension was then layered over 3ml FBS, centrifuged for 10 min at 300g. The pellet was resuspended in 90% FBS-10% DMSO, 5 million cells/mL, and cells stored in liquid nitrogen.

#### Treg and Tconv magnetic isolation

Frozen PBMCs samples were processed in batches of 4 to 10 samples, with at least one HD by batch, with strict attention to processing time to avoid cell aggregation, which was otherwise pervasive with samples from severe COVID-19 patients. 5-10×10^5^ PBMCs were resuspended into PBS with 2% FBS and 1mM EDTA. CD4+CD25hiCD127low (Treg) were isolated by positive and negative magnetic selection, CD4+CD25-(Tconv) by negative selection only. EasySep™ Human CD4+CD127lowCD25+ Regulatory T Cell Isolation Kit (StemCell, #18063) was used following manufacturer’s instructions. 10% of the final isolated fraction was stained with anti-CD4, anti-CD8, anti-CD14, anti-CD19 and live dead for 15 min at 4°C, fixed in 1% PFA for 10 min at room temperature and analyzed by flow cytometry to determine purity and yield (antibodies references below). The remaining 90% of the samples were resuspended in lysis buffer (TCL Buffer (QIAGEN) supplemented with 1% 2-Mercaptoethanol) at an average concentration of 500-1,500 cells per 5ul, and stored in a low-binding tube at −80°C.

#### Flow cytometry

Cells were first incubated in 100 μL of PBS with 2mM EDTA for 15 min with 5μL Fc Block (Human TruStain FcX™, Biolegend, cat #422301) and a 1:500 dilution of Zombie UV viability dye (Biolegend, cat# 423107). They were then washed with FACS buffer (phenol red–free DMEM, 2% FBS, 2mM EDTA, 10mM HEPES),and stained at 4°C for 25 minutes using the following cell surface antibodies: CD14 Pacific Blue (clone M5E2, BioLegend cat# 301815, 2:100 dilution); CD19 Pacific Blue (clone H1B19, BioLegend cat#302224, 2:100 dilution); CD3 AF700 (clone OKT3, BioLegend, cat# 317340, 2:100 dilution); CD4 PerCP-Cy5.5 (clone OKT4, BioLegend, cat# 317428, 2:100 dilution); CD127 AF488 (clone A019D5, BioLegend, cat# 351314, dilution 3:100); CD25 PE-Cy7 (clone BC96, BioLegend, cat# 302611, dilution 3:100); KLRG1 APC-Cy7 (clone 2FI/KLRG1, BioLegend, cat# 138426, 2:100 dilution); CD279 (PD-1) BV650 (clone EH12.2H7, BioLegend, cat# 329950, dilution 2:100); CD45RA BV510 (clone HI100, BioLegend, cat# 304142, dilution 2:100). After cell surface staining, cells were fixed overnight at 4°C using 100 μL of Fix/Perm buffer (eBioscience), followed by permeabilization using 1X permeabilization buffer (eBioscience) for 40 minutes at room temperature in the presence of the following intracellular antibodies: FoxP3 APC (clone PCH101, Invitrogen, cat# 17-4776-42, dilution 4:100); Tbet BV605 (clone 4B10, BioLegend, cat# 644817, dilution 4:100); Bcl6 PE/Dazzle (clone 7D1, BioLegend, cat# 358510, dilution 4:100). Data was recorded on a FACSymphony^TM^ flow cytometer (BD Biosciences) and analysed using FlowJo 10 software.

#### RNAseq

##### Low-input RNAseq

RNAseq was performed in two different batches, including samples coming from 2 to 7 different experimental batches (different experiment dates in Table S1). After the magnetic isolation and using the data from the flow cytometry post-isolation, samples with purity > 65% and expected number of cells >800 were selected for RNAseq. RNA-seq was performed with 5μL of the previously described lysate following the standard ImmGen low-input protocol (www.immgen.org). Smart-seq2 libraries were prepared as previously described (Picelli et al., 2014) with slight modifications. Briefly, total RNA was captured and purified on RNAClean XP beads (Beckman Coulter). Polyadenylated mRNA was then selected using an anchored oligo(dT) primer (50–AAGCAGTGGTATCAACGCAGAGTACT30VN-30) and converted to cDNA via reverse transcription. First strand cDNA was subjected to limited PCR amplification followed by Tn5 transposon-based fragmentation using the Nextera XT DNA Library Preparation Kit (Illumina). Samples were then PCR amplified for 12 cycles using barcoded primers such that each sample carries a specific combination of eight base Illumina P5 and P7 barcodes for subsequent pooling and sequencing. Paired-end sequencing was performed on an Illumina NextSeq 500 using 2 x 38bp reads with no further trimming. Reads were aligned to the human genome (GENCODE GRCh38 primary assembly and gene annotations v27) with STAR 2.5.4a (https://github.com/alexdobin/STAR/releases). The ribosomal RNA gene annotations were removed from GTF (General Transfer Format) file. The gene-level quantification was calculated by featureCounts (http://subread.sourceforge.net/). Raw read counts tables were normalized by median of ratios method with DESeq2 package from Bioconductor (https://bioconductor.org/packages/release/bioc/html/DESeq2.html) and then converted to GCT and CLS format.

##### Quality control

Samples with less than 1 million uniquely mapped reads were automatically excluded from normalization to mitigate the effect of poor-quality samples on normalized counts. Samples having fewer than 8,000 genes with over ten reads were also removed from the data. We screened for contamination by using known cell type specific transcripts (per ImmGen ULI RNAseq and microarray data). Finally, the RNA integrity for all samples were measured by median TIN across human housekeeping genes with RSeQC software (http://rseqc.sourceforge.net/#tin-py). Samples with TIN < 45 were removed from the data set prior to downstream analysis.

##### Batch correction

In order to analyze simultaneously the two RNAseq batches, the read counts tables from these two batches were first combined for all the Treg samples and for all the Tconv samples separately. Then, the batch effects were corrected by using Combat method from sva package.

##### Viral reads mapping

The SARS-Cov-2 genome sequence and annotation were obtained from NCBI (https://www.ncbi.nlm.nih.gov/sars-cov-2/). Reads were aligned by STAR 2.5.4a with the parameters suggested in Kim et al.(Kim et al., 2020). The gene-level quantification was calculated by featureCounts (Subread 1.6.2). Read counts tables were normalized by median of ratios method with DESeq2 package (Love et al., 2014)

##### Regression Linear model

Related to the epidemiology of the disease, there was a strong sex-bias in severe patients in our dataset (predominantly male). Thus, we performed a linear regression (*glm()* function in R) using log-transformed expression as the response variable, and severity and sex as explanatory variables. We removed 45 genes highly correlated with sex, which were no longer associated with severity once adjusted.

##### Differential gene expression

After quality control and regression, genes with a minimum reads count of 20 in more than 20% of samples from a population (Treg or Tconv) were retained. We used an uncorrected t-test to compute differential gene expression between the different groups from the normalized read counts dataset. Genes with a FoldChange >2 or <0.5 and p-value < 0.05 were selected for further analysis.

##### Computation of signatures and module indexes

The different COVID-19 (SCTS), Treg signature, TITR, and modules indexes were calculated for each donor by averaging the normalized expression (versus mean of all HD) of all genes belonging to each signature.

##### Geneset enrichment analysis

CD4+ Tcell gene signatures were curated from published and relevant datasets, as described (Zemmour et al., 2018). Only datasets containing replicates were used. To reduce noise, genes with a coefficient of variation between biological replicates <0.6-0.8 in either comparison groups were selected. Up- and downregulated transcripts were defined as having a fold change in gene expression >1.5 or <0.6 and a t-test p-value <0.05, limited to the top 300 genes by signature. Other signatures were obtained from extracting all CD4+ T cells signatures from the MSigDB C7 Immunologic collection (Godec et al., 2016). Geneset enrichment analysis with the COVID-19 signature was performed using hypergeometric distribution and type I error was controlled using FDR. Signatures with FDR <10% and an overlap with COVID-19 signature >10 genes were considered as significant and are reported in table S3.

#### scRNAseq re-analysis

Data deposited at data repository cellxgene http://doi.org/10.5281/zenodo.3710410 were used, with the cell annotation provided by Wilk *et al* (Wilk et al., 2020), CD4+ and CD8+ cell clusters were extracted from the processed single cell data. Using the Seurat pipeline (Butler et al., 2018), principal components and UMAP coordinates were recomputed for just the CD4+ and CD8+ T cell populations. From there, K-means clustering was re-computed using the FindClusters function with default parameters and the cluster with the highest average expression for a list of core Treg signature genes (*CAPG, FOXP3, TNFRSF4, IFI27L2A, TNFRSF18, FOLR4, TNFRSF9, S100A6, APOBEC3, IKZF2, H2AFZ, CTLA4, LY6A, HOPX, SERINC3,* and *IL2RA;* from https://www.biorxiv.org/content/10.1101/2020.07.06.189589v1) was flagged for further analysis. From this Treg cluster, cell averages were calculated across all genes of the Severe COVID-19 (SCTS) UP signature, and color-coded on the Treg UMAP space for Fig. 3H.

#### Statistics

Unless specified otherwise, data are presented as mean ± SD and tests of associations for different variables between COVID-19 patients and HD were computed (1) using the nonparametric Mann Whitney test or (2) using randomization test: random values were generated (nrorm() in R) from the mean and standard deviation of log-transformed values in HD controls, testing the frequency of draws that led to a number of observations > 95^th^ quantile of HD values that was equal or greater to the number of such observations in each patient group. Correlation coefficients were from Pearson correlation. Significance of signature overlaps into our dataset was assessed by Fisher’s exact test when computing one signature at a time, or by a hypergeometric test with Benjamin–Hochberg correction when using the large curated CD4+ T cell signatures database. Analyses and plots were done using RStudio® (v.1.2.5019) and GraphPad Prism® (v.8.4.3), heatmaps generated with Morpheus (https://software.broadinstitute.org/morpheus).

#### Data availability

The data reported in this paper have been deposited in the Gene Expression Omnibus (GEO) database under accession no. GSEXXX (human RNAseq).

## SUPPLEMENTARY FIGURE LEGENDS

**Fig. S1 (related to Fig. 1).**
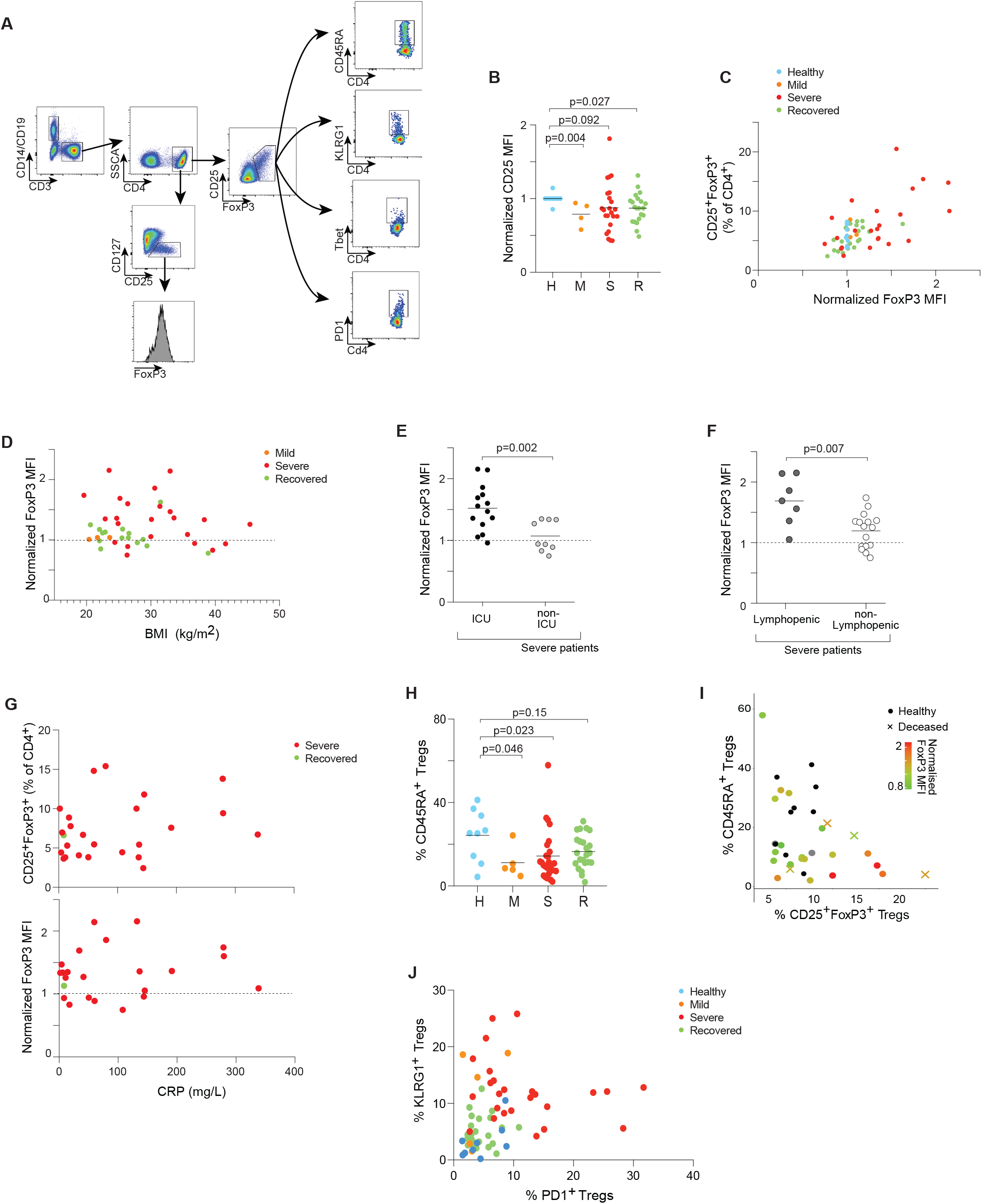
Flow cytometry phenotypes of Tregs from severe COVID19 patients. (A) Representative flow cytometry plots of the gating strategy for Tregs, FoxP3 MFI and Treg markers. (B) Expression of CD25, measured as normalized MFI in CD127^lo^ CD25^+^ Tregs across the whole cohort. (C) Correlation between percentage of CD25^+^ FoxP3^+^ Tregs and FoxP3 expression as MFI across the whole cohort. (D) Correlation between FoxP3 expression as MFI in Tregs and BMI across the whole cohort. (E,F) Expression of FoxP3 as MFI in Tregs from severe COVID-19 patients, comparing intensive care unit (ICU) admission (G) and lymphopenia, determined as a B cell or T cell percentage significantly lower than the average (D). (H) Correlation between CRP level in COVID patients and percentage of Tregs (top) or expression of FoxP3 as MFI (bottom). (I) Proportion of CD45RA+ Tregs as determined by flow cytometry across the whole cohort; p-values from Mann–Whitney test. (J) Correlation between percentage of CD25^+^ FoxP3^+^ Tregs (x-axis), percentage of CD45RA+ Tregs (y-axis) and FoxP3 expression as MFI (color gradient) within severe COVID-19 patients. Healthy controls depicted in black dots and patients with fatal outcome by a cross. (K) Correlation between KLRG1+ Tregs and PD1+ Tregs as determined by flow cytometry, across the whole cohort. All P-values were computed from Mann–Whitney test.

**Fig. S2 (related to Fig. 2).**
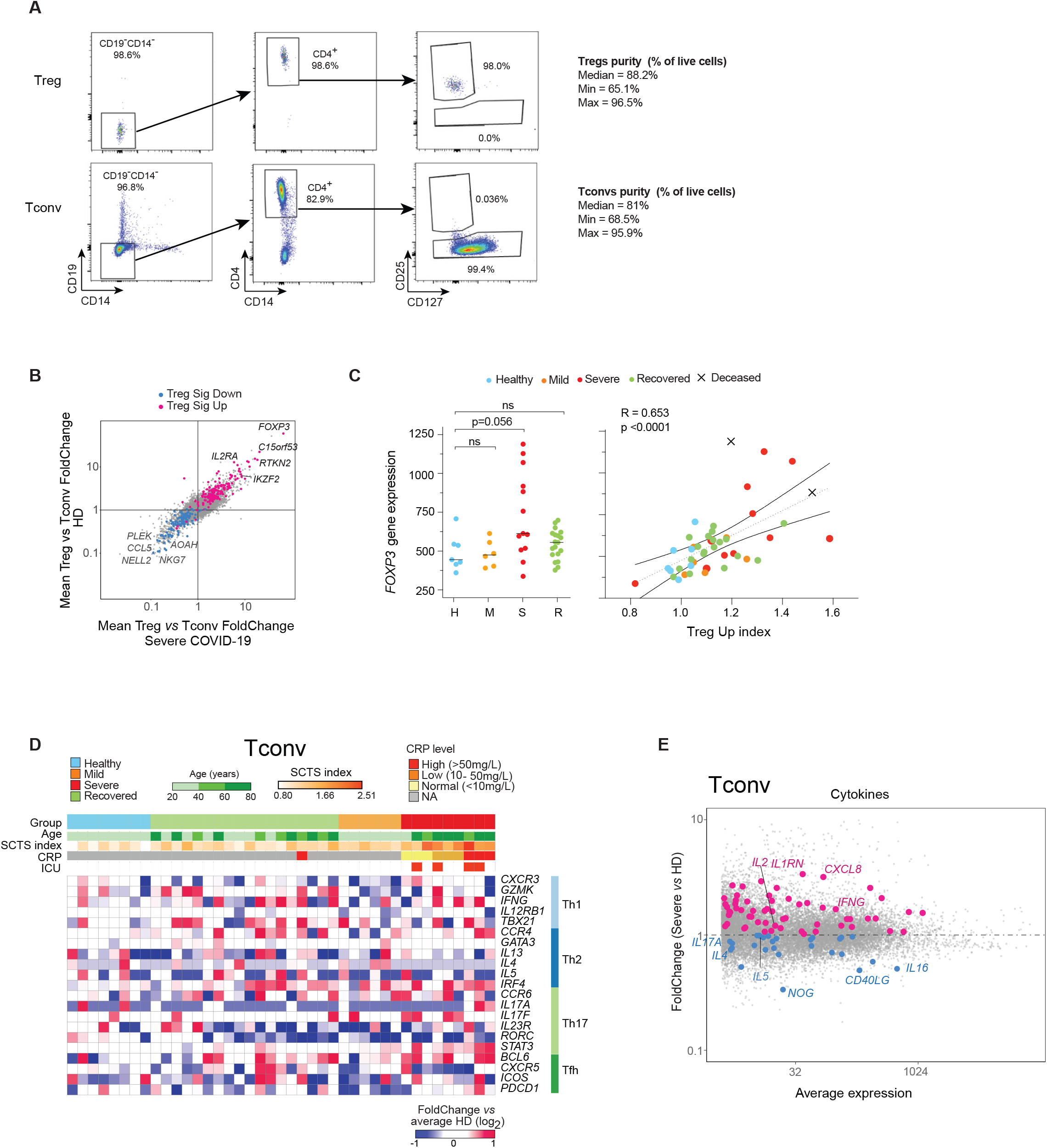
Transcriptomic profile of CD4+ Treg and Tconv cells. (A) Representative flow cytometry plots showing a sample of the final isolated fractions of magnetically-purified purified Tregs (top) and Tconvs (bottom). The median and min/max of purity for each isolated population in the whole cohort is listed. (B) Comparison between the Treg/Tconv FoldChange in HD (y-axis) versus severe COVID-19 (x-axis); Treg signature genes (Ferraro et al., 2014) are highlighted. (C) Gene expression of *FOXP3* in Tregs across the whole cohort (left) and its correlation with the TregUP index (right). Patients with fatal outcome depicted by a cross. R statistic from a Pearson correlation procedure and p-values from Mann–Whitney test. (D) Expression heatmap of the canonical T helpers’ genes (Th1, Th2, Th17 and T follicular helpers) in Tconvs from each patient versus average expression in HD. One column per patient, with severity groups color-coded, and with color-gradients for age, *Severe COVID19 Treg Signature* (SCTS) index and systemic level of CRP. (E) FoldChange vs average expression (MA) plot from severe COVID-19 patients’ Tconvs compared to HD. Upregulated (pink) and downregulated (blue) cytokine transcripts highlighted.

**Fig. S3 (related to Fig. 3).**
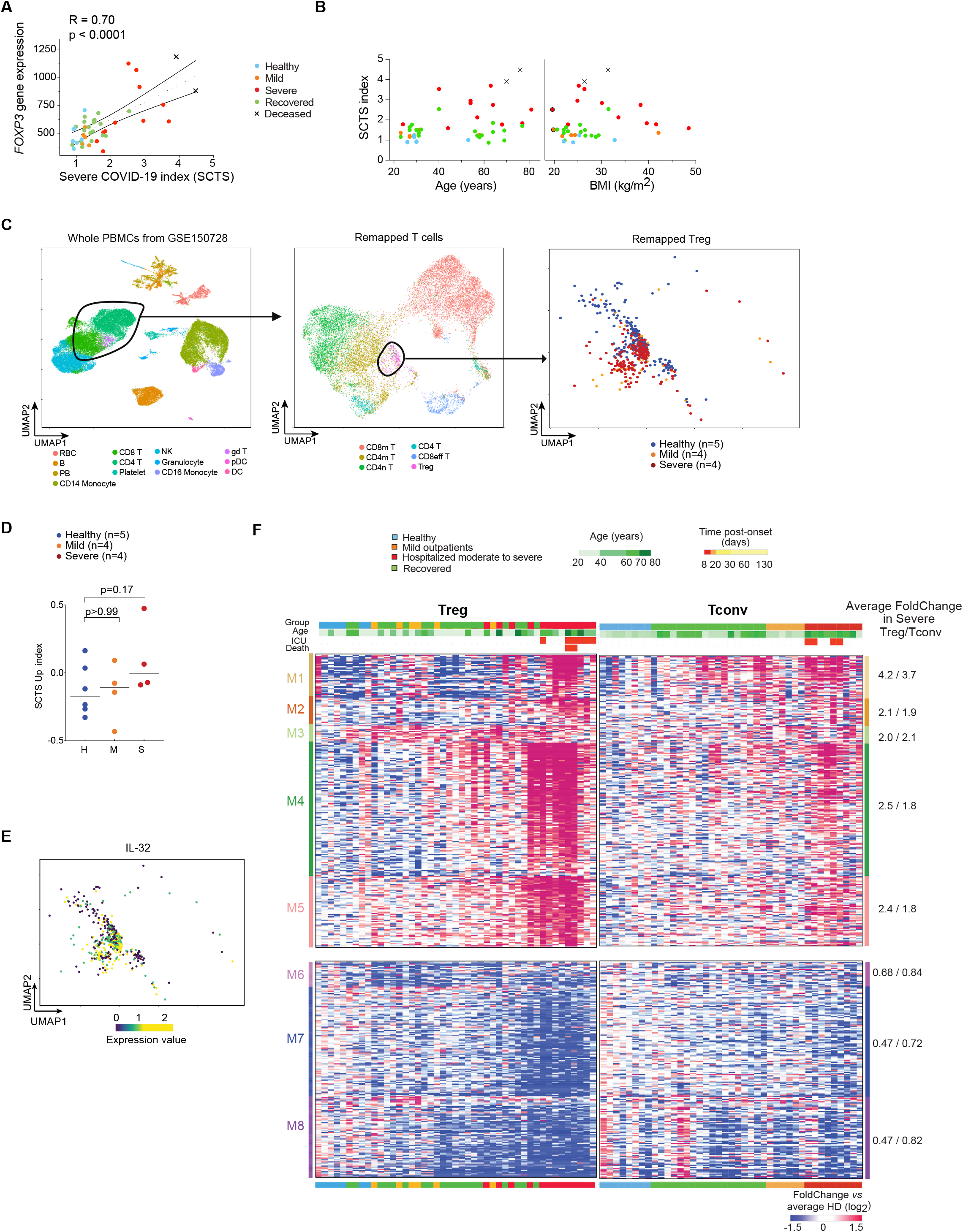
Elements of the severe COVID-19 Treg signature. (A) Correlation between *FOXP3* gene expression and the *Severe COVID19 Treg Signature* (SCTS) index across the whole cohort. (B) Correlation between SCTS index and age (left) or BMI (right) across the whole cohort. Patients with fatal outcome depicted by a cross. R statistic and p-value from a Pearson correlation procedure. C) Extraction strategy of the Tregs population from scRNAseq dataset (GSE150728). COVID-19 patients’ PBMCs were displayed as a 2D UMAP at different levels of extraction: the whole PBMCs (left panel), T cells (middle panel) and Tregs (right panel). Samples are color-coded by cell type, except for Tregs which are color-coded by patient group. (D) Quantification of the SCTS index for each patient among the different groupsp-values from Mann–Whitney test. (E) Same 2D UMAP than in A (right panel) but indicating *IL32* expression among the COVID-19 Tregs dataset. (F) Heatmap of the differentially expressed genes identified in Fig 3B (p<0.05 (t-test), FC >2 or <0.5) across all groups in Tregs (left) versus Tconvs (right). Each column represents one sample. Top ribbons indicate for each individual: severity group, age, ICU admission and final outcome (deceased, in red). Left ribbon indicates the different modules and the right ribbon the average FoldChange in severe Tregs versus severe Tconvs.

**Fig. S4 (related to Fig. 4).**
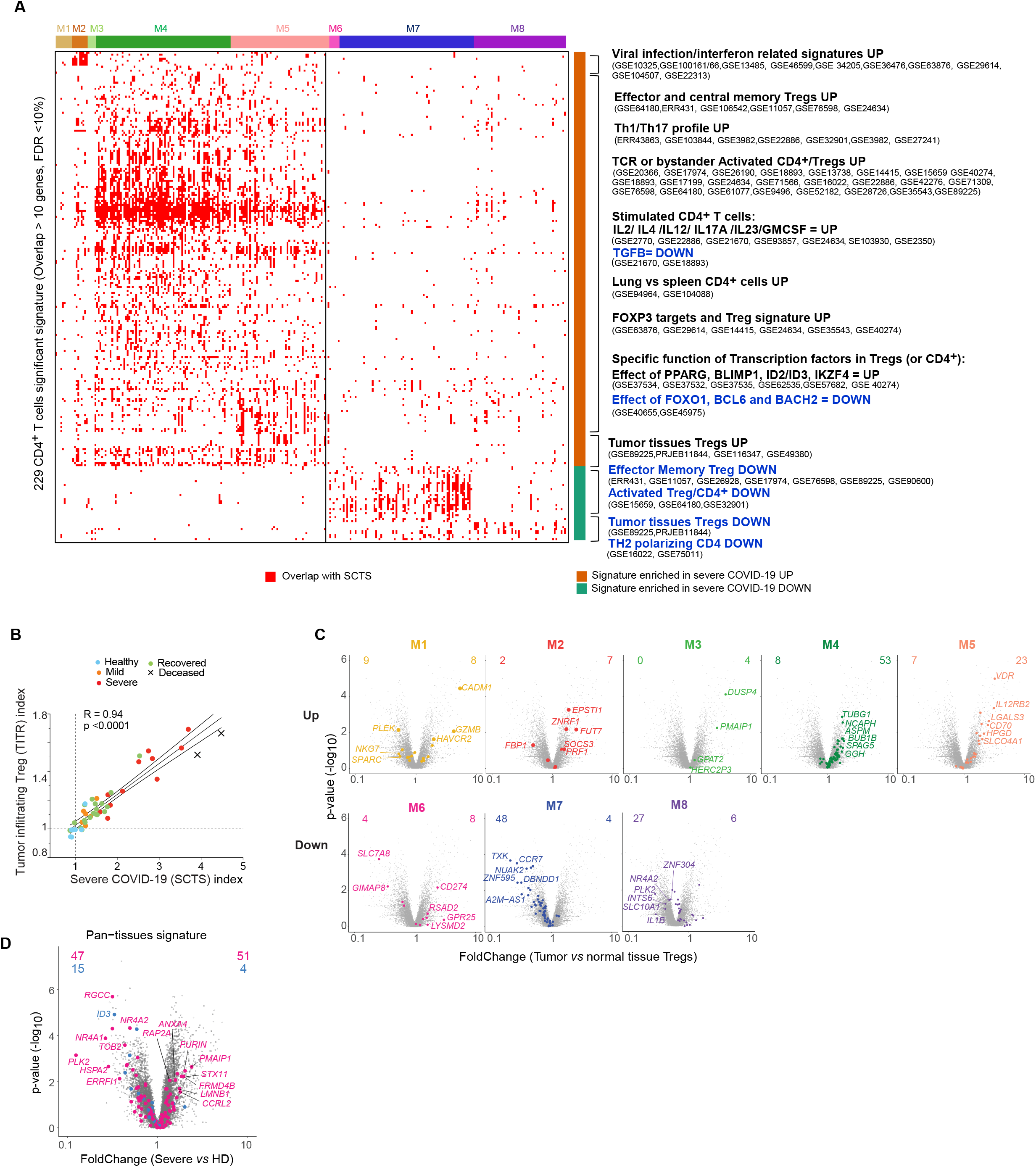
Similarities between tumor and severe COVID-19 Treg. (A) Heatmap of the overlap between the *Severe COVID19 Treg Signature* (SCTS) (defined in Fig3A), and other relevant immunological signatures (hypergeometric test FDR <10% and more than 10 genes in overlap). The top ribbon indicates the modules (defined in Fig3C). Annotation of common clusters of signatures and the signatures belonging to these clusters with their GEO dataset ID at the right. All signatures and the overlap parameters can be found in Table S3. (B) Correlation between the TITR (Tumors infiltrating Tregs index) and the SCTS index. Patients with fatal outcome depicted by a cross. R statistic and p-value from Pearson correlation procedure. (C) FoldChange vs p value (volcano) plots of normalized expression in Tregs from colorectal cancer (CRC) vs normal colon Tregs (Magnuson et al., 2018). SCTS modules from Fig 3C are highlighted and the numbers of their genes up and down are annotated. (D) FoldChange vs p value (volcano) plot of normalized expression in Tregs from severe COVID-19 patients compared to HD. Signature genes from Pan Tissues Tregs (Dispirito et al., 2018) are highlighted.

## SUPPLEMENTARY TABLE LEGENDS

**Table S1. Clinical, biological and analytic characteristics of the cohort (n=68)**

**(A)**Clinical characteristics of COVID-19 patients and healthy donors (summary, n=68) **(B)**Individual clinical and biological data (n=68) - at collection date **(C)**Samples included in the Flow cytometry dataset (n=63): key parameters **(D)**Samples included in the Low input RNAseq dataset: purity and quality (Tregs n=45 & Tconvs n=41) **(E)**Viral reads in the Treg/Tconv RNAseq dataset. NA: Not available.

**Table S2. *Severe COVID19 Treg Signature* (SCTS) genes with their average Severe/HD fold change in Tregs**

**Table S3. CD4+ signatures significantly enriched in the *Severe COVID19 Treg Signature* (hypergeometric test FDR <10% and more than 10 genes in overlap).**

